# TSEBRA: Transcript Selector for BRAKER

**DOI:** 10.1101/2021.06.07.447316

**Authors:** Lars Gabriel, Katharina J. Hoff, Tomáš Brůna, Mark Borodovsky, Mario Stanke

## Abstract

**Background:** BRAKER is a suite of automatic pipelines, BRAKER1 and BRAKER2, for the accurate annotation of protein-coding genes in eukaryotic genomes. Each pipeline trains statistical models of protein-coding genes based on provided evidence and, then predicts protein-coding genes in genomic sequences using both the extrinsic evidence and statistical models. For training and prediction, BRAKER1 and BRAKER2 incorporate complementary extrinsic evidence: BRAKER1 uses only RNA-seq data while BRAKER2 uses only a database of cross-species proteins. The BRAKER suite has so far not been able to reliably exceed the accuracy of BRAKER1 and BRAKER2 when incorporating both types of evidence simultaneously. Currently, for a novel genome project where both RNA-seq and protein data are available, the best option is to run both pipelines independently, and to pick one, likely better output. Therefore, one or another type of the extrinsic evidence would remain unexploited.

**Results:** We present TSEBRA, a software that selects gene predictions (transcripts) from the sets generated by BRAKER1 and BRAKER2. TSEBRA uses a set of rules to compare scores of overlapping transcripts based on their support by RNA-seq and homologous protein evidence. We show in computational experiments on genomes of 11 species that TSEBRA achieves higher accuracy than either BRAKER1 or BRAKER2 running alone and that TSEBRA compares favorably with the combiner tool EVidenceModeler.

**Conclusion:** TSEBRA is an easy-to-use and fast software tool. It can be used in concert with the BRAKER pipeline to generate a gene prediction set supported by both RNA-seq and homologous protein evidence.

## Background

Currently, the National Center for Biotechnology Information’s (NCBI) GenBank [1] hosts 7,978 eukaryotic genomes, with 3,208 of these genomes lacking an annotation of protein-coding genes. Notably, 746 genome annotations out of existing 4,770 ones were generated by NCBI [2, 3]. The original authors frequently omit an annotation step and many publicly available genomes remain not annotated. Furthermore, re-annotation may be in order for many of the annotated genomes as more related sequence data has become available, or annotation methods have been improved since their initial application. Thus, there is a need for accurate automated annotation methods that use all available data and are easily accessible to bioinformatics teams.

The most useful data to support accurate genome annotation are transcriptomic sequence data, e.g. RNA-sequencing (RNA-seq) data, from the same species and protein sequences from species that are sufficiently closely related to the target species in the tree of life. RNA-seq reads spliced aligned to a genomic region are used to infer likely intron intervals [4] in protein-coding genes. In a similar way, likely exon and intron boundaries can be inferred using homologous proteins because segments of gene structures are often highly conserved [5]. Protein evidence has the advantage that it maps only to protein-coding genes, but with the downside that it depends on the degree of sequence conservation, which may differ between genes and available species. In contrast, RNA-seq is usually obtained from the same species and then, free of this dependency, covers only genes and spliced isoforms that are expressed in a sample. However, RNA-seq could be generated from non-coding genes; sequencing errors may render accurate alignments difficult. Ever-increasing throughput has resulted in large databases of RNA-seq. For example, the NCBI Sequence Read Archive (SRA) [6] hosts more than 36 petabytes of data, while the protein database OrthoDB [7] contains more than 37 million sequences.

Genome annotation methods that use statistical models of gene structures such as splice site patterns in addition to the evidence from RNA-seq and homology, are arguably best suited for whole-genome annotation [8]. BRAKER, a popular pipeline of competitive accuracy [9], has two modes of a genome annotation process supported by extrinsic evidence. BRAKER1 uses GeneMark-ET [10–12] together with AUGUSTUS [13–17] and relies on RNA-seq data to support gene finder training and accurate prediction of gene structures. BRAKER2 [18] exploits spliced alignments of homologous proteins as a source of extrinsic evidence for genome annotation with GeneMark-EP+ [19] and AUGUSTUS.

When heterogeneous extrinsic evidence sources are available, some genome annotation tools like MAKER2 [20] and GeMoMa [21] integrate these different sources directly into the annotation protocol. Some, like the recent FINDER [22], perform protein-spliced alignments only with proteins that are mapped to genes missed by RNA-seq-based methods. On the other hand, FINDER does not use RNA-seq evidence to assess or compare homology-based gene models. A different approach is to first generate multiple whole-genome annotations and then to use a *combiner tool* that takes various gene predictions as input with diverse sources of extrinsic evidence and constructs a genome annotation that is on average more accurate than any input genome annotation. Some previously developed combiner tools built their own gene structure model in the form of a graph and report a gene structure either based on the consensus of all available data, e.g. IPred [23], or as the result of a machine learning procedure such as most likely parse of an HMM, e.g. Combiner [24], JIGSAW [25], Evigan [26], ExonHunter [27]. A prominent combiner tool is the openly accessible EVidenceModeler (EVM) [28]. It uses a weighted consensus from all available evidence sources to predict a gene structure. EVM was successfully used to produce several high-quality annotations of novel genomes [29, 30].

In our approach, we first generate several sets of whole-genome gene predictions based on a single type of extrinsic evidence (i.e. by BRAKER1 and BRAKER2). We use a new combiner tool that scores and ranks these predictions (transcripts) based on heterogeneous evidence. Then, we select those with higher rank into a newly constructed genome annotation which is on average more accurate than any whole-genome annotation provided in the input. Up to now, the BRAKER suite has so far not been able to achieve a prediction accuracy that is reliably superior to either single-source evidence mode when using RNA-seq and proteins simultaneously [31, 32]. Nevertheless, BRAKER users often have both types of extrinsic data available for a target genome. Incidentally, most of the previously mentioned combiner tools are either not publicly available anymore, lack support, or are very difficult to use for combining BRAKER1 and BRAKER2 predictions. Therefore, we present *Transcript Selector for BRAKER (TSEBRA)*, a fast software tool for selecting gene predictions from the output of two branches of the BRAKER eukaryotic gene prediction suite based on all the heterogeneous extrinsic evidence. TSEBRA achieves high accuracy and is easy to use. We show that it delivers a significant increase in accuracy with respect to the input annotations generated by BRAKER1 and BRAKER2.

## Implementation

TSEBRA uses a set of arbitrarily many gene prediction files in GTF format together with a set of files of heterogeneous extrinsic evidence to produce a combined output. From the whole set of transcripts contained in the gene predictions, TSEBRA must select those that are more reliably supported by a full complement of extrinsic evidence; these transcripts constitute the output. Less reliably supported transcripts are filtered out. The rational of TSEBRA’s approach is as follows. Taking a union of gene predictions generated by two or more gene finding tools makes a set of predictions with improved sensitivity but with lower specificity. A non-trivial task is to remove some predictions and increase specificity with little decrease of sensitivity. This task is tantamount to identification of likely false positives and filtering them out. TSEBRA solves exactly this problem.

TSEBRA uses extrinsic evidence in the form of intron regions or start/stop codon positions to evaluate and filter transcripts from gene predictions. These must be provided in a GFF file that includes two attributes in the last column ‘mult=’, a number specifying its *multiplicity* – the number of alignments that support it, and ‘src=’ determining its source, e.g., ‘src=P’ for evidence from a protein alignment. The mult attribute is used to specify multiplicities larger than one.

TSEBRA takes three sets of different hyperparameters from a configuration file. More precisely, it takes a weight for any evidence source, four transcript score thresholds and two low evidence support thresholds. The weights are used to compute *transcript scores* and the transcript score thresholds are used for comparing transcripts. The low evidence support thresholds consist of minimum fractions of intron or start/stop codon support. We recommend the application of the default hyperparameters provided in the TSEBRA configuration file to be used in a standard use case.

The workflow of TSEBRA is as follows:

1. Take a union of transcripts predicted by BRAKER1 and BRAKER2 while merging identical transcripts.
2. Compute vectors of support scores for all transcripts.
3. Identify all pairs of transcripts with overlapping coding regions.
4. Compare all pairs of overlapping transcripts by a transcript comparison rule using the extrinsic evidence and mark some of them for exclusion.
5. Remove all transcripts marked for exclusion by the transcript comparison rule.
6. Remove all transcripts with low evidence support.
7. Combine the remaining transcripts into a final set of predictions with groups of overlapping transcripts making sets of alternative isoforms.

The output of TSEBRA is the set of genes (with alternative isoforms) in GTF format.

In step 6, a transcript is removed if the fractions of introns and start/stop codons supported by extrinsic evidence are lower than the low evidence support thresholds. In step 7, genes are the single-linkage clusters of transcripts where two transcripts are in the same gene if they overlap (and could be alternative splice forms). Two transcripts are considered to overlap if they share at least three adjacent protein-coding nucleotides on the same strand and in the same reading frame. Note that a transcript ‘marked for exclusion’ in step 4 is still compared to all overlapping transcripts and may cause removal of another transcript. This filtering step is different from a simplistic approach that would first score transcripts and then apply a fixed threshold to their score. In our approach, the transcripts with strongest local support are kept, and those that are discarded can still have strong support in absolute terms if transcripts with even stronger support overlap.

As a special case, TSEBRA may be used with a single gene prediction file to filter for the ones with the strongest evidence support. This may be useful for a genome annotation with many transcript isoforms per gene.

### Transcript scores

Four transcript scores *s*_1_, …, *s*_4_ characterize the support of *features* of a transcript, here introns (i) or start/stop-codons (s), by all extrinsic evidence *E* represented by *hints*. A hint *h* is either an intron region or start/stop codon position together with an identifier of its original source *src*(*h*) ∈ *O* and its multiplicity *mult* (*h*) ∈ {1, 2 *…*}. *O* is a set of original sources, e.g. *O* = {P, R}, if protein data and RNA-seq were used, but could also contain further elements, e.g. when variants of RNA-seq sequencing technologies shall be distinguished [32]. Multiplicity *mult* (*h*) is the number of alignments from the same source that supports hint *h*. A hint supports a transcript feature if all identifying characteristics match, i.e. sequence name, start/stop position, feature type, and strand.

Consider a particular transcript and let *F* be the set of all of its features. Define *F_f_* ⊂ *F* as all features in *F* of type *f* ∈ {i, s}. The relative support of a transcript feature is

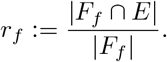

Score *s*_1_ := *r*_i_ is the relative support of the transcript’s introns (*f* = i) by the evidence *E* and *s*_2_ := *r*_s_ is the fraction of start/stop codons supported by *E*.

A weight *w_o_* with *o* ∈ *O* is assigned to each evidence source. The absolute quantity of supporting hints for a transcript feature *f* is the weighted sum of all supporting hints:

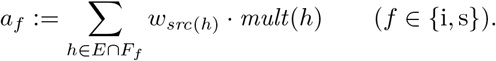

The scores *s*_3_ := *a_i_* and *s*_4_ := *a_s_* measure the abundance of extrinsic evidence that support the introns or the start/stop codons of a transcript, see Figure 1 for an example.

**Figure 1.**
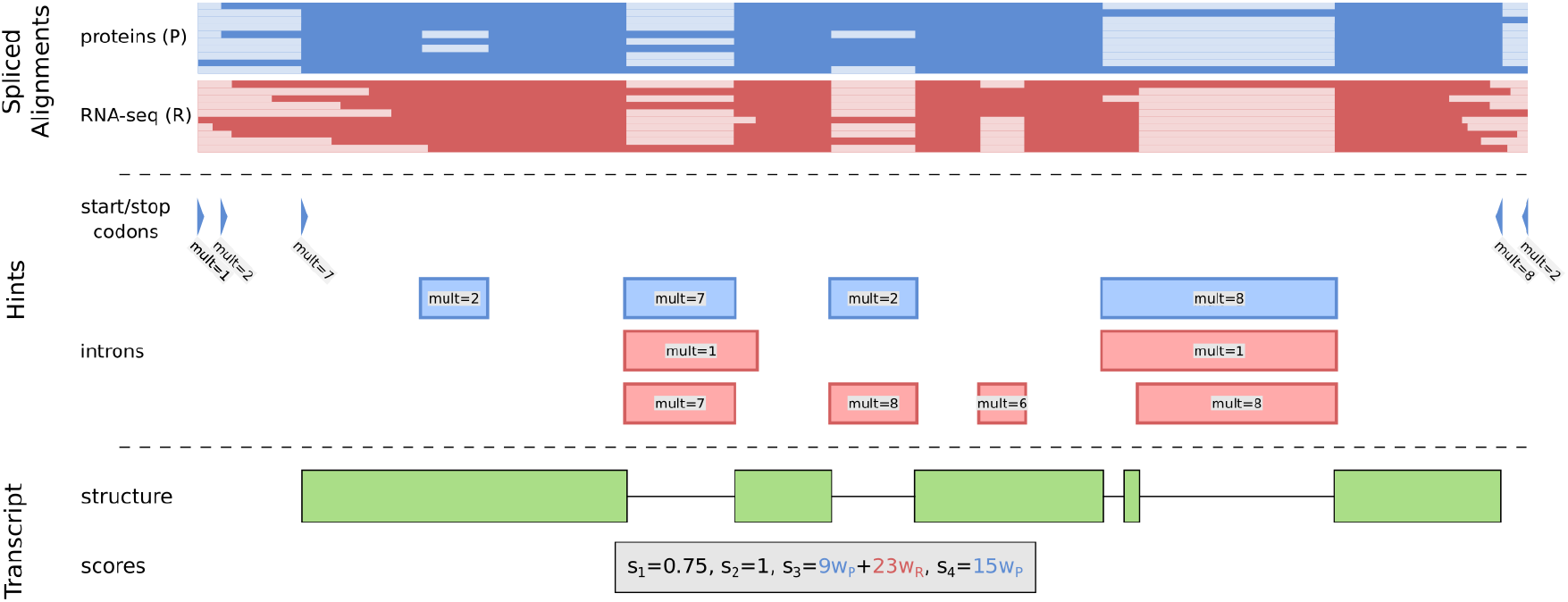
Example of how extrinsic evidence in form of spliced alignments from homologous proteins (blue) or RNA-seq reads (red) is used to determine scores for the support of a transcript (green). Likely exon-intron borders are inferred from the alignments to create intron hints. The start and stop codons of the protein alignments are used to create start and stop codon hints, respectively. The transcript scores utilize them to quantify the support of the transcript structure.

### Pairwise transcript comparison rule

The pairwise transcript comparison rule compares two transcripts with respect to their support of extrinsic evidence using the transcript scores. One or no transcript is marked for exclusion when comparing two overlapping transcripts, see Figure 2. The differences of all transcript scores (of the same type) are compared to a score specific threshold, in order from *s*_1_ to *s*_4_. When the threshold is exceeded for the first time, the comparison rule terminates and the transcript with the smaller value for the current score is marked as the transcript that will be excluded from the combined gene set. Neither transcript is marked for removal if all differences are less than or equal to the associated thresholds.

**Figure 2.**
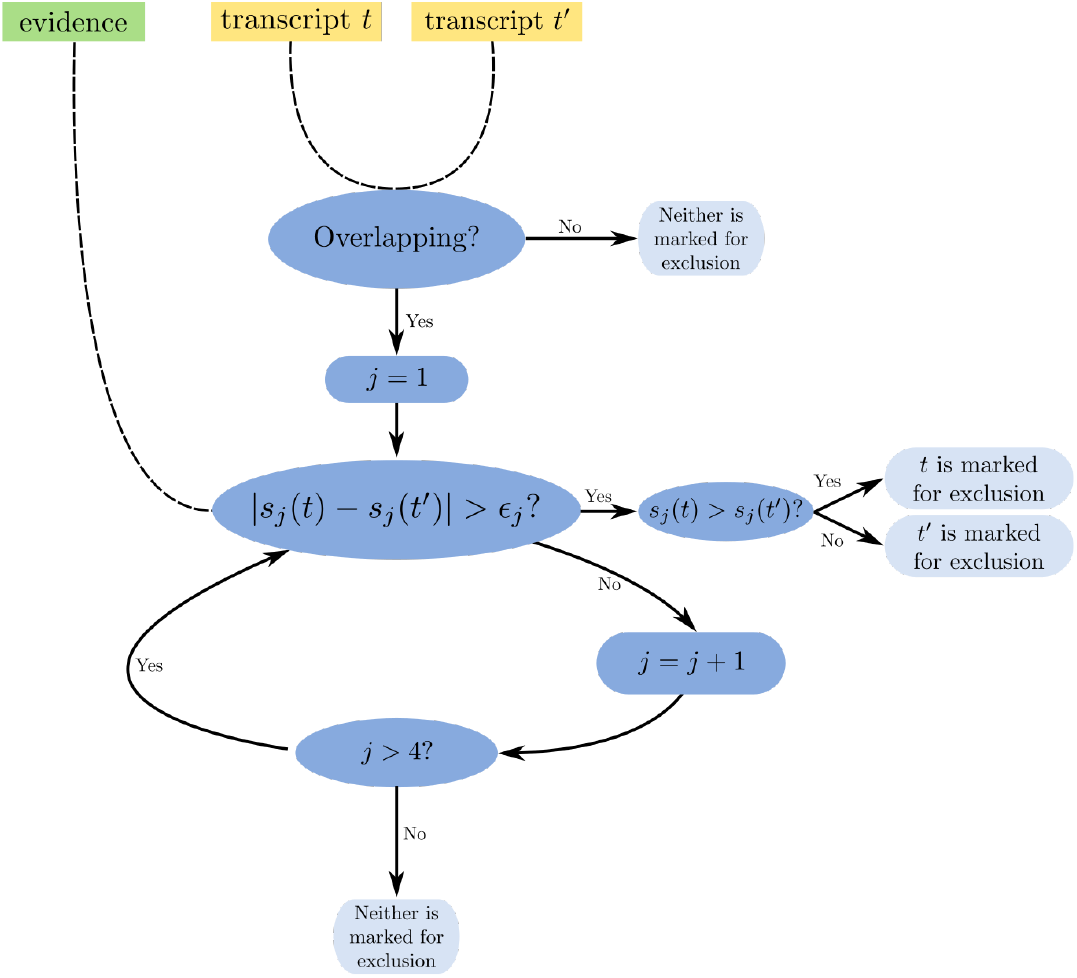
Comparison rule for two transcripts using extrinsic evidence, either one or none of the transcripts is marked for removal; *s_j_* are transcript scores and ϵ*_j_* are score specific thresholds with *j* ∈ {1, 2, …, 4}.

### Default hyperparameters

The TSEBRA suite includes a set of default hyperparameters, which are recommended for usage in a standard use case – to combine BRAKER1 and BRAKER2 – so that users are not required to set the hyperparameters themselves. Evidence sources in a standard BRAKER1 and BRAKER2 output are: protein database (P), EST database (E), combined EST/protein database (C), and manual anchored (M). The default weights for these are *w_P_* = 0.1, *w_E_* = 10, *w_C_* = 5 and *w_M_* = 1. A transcript has low evidence support in this default setting if the fraction of supported introns is less than 0.75 and the supported start/stop-codon fraction is less than 1.0. The score specific thresholds are *ϵ*_1_ = 0, *ϵ*_2_ = 0.5, *ϵ*_3_ = 25, *ϵ*_4_ = 10. We have shown that TSEBRA using default parameters performs with high accuracy across several species, see Results and discussion.

## Results and discussion

We compared the accuracy of TSEBRA in two experiments. First, we compared TSEBRA to BRAKER1 and BRAKER2 in their standard use modes, and second, we compared TSEBRA with EVM.

### Accuracy assessment metrics

Specificity (Sp), sensitivity (Sn), and their harmonic mean – the F1-score – were the measures of gene prediction accuracy. Accuracy values were examined at the gene, transcript, and exon levels. A predicted gene is considered correct, if it is identical to at least one annotated alternative splicing isoform. A reference transcript *t* is considered as correctly predicted by transcript *t′*, if *t* and *t′* completely agree on their sets of CDS (exons). Two CDS are considered to agree if they are located in the same strand and both pairs of sequence coordinates are identical.

### Comparison with BRAKER1 and BRAKER2

Complete genome annotations generated independently by BRAKER1 and BRAKER2 (both BRAKER v.2.1.5) for 11 eukaryotic species (Table S1 in Supplementary Information) were processed by TSEBRA with default hyperparameters. For each genome, we used its ‘standard’ annotation to compute the accuracies of the sets of gene predictions made by BRAKER1, BRAKER2, and TSEBRA. BRAKER1 was supported by extrinsic evidence in form of RNA-seq reads aligned to the genome of interest. RNA-seq hints were sampled with VARUS [33] from SRA for each genome with HISAT2 [34] as an alignment tool. BRAKER2 was supported by protein data sets selected earlier while testing BRAKER2 [18]. For each of the three model species with genome annotations curated multiple times, *A. thaliana*, *C. elegans*, and *D. melanogaster*, we used proteins from three sets of species varied with respect to minimal evolutionary distance to the query species. Each protein set included proteins from a large clade, e.g. Plantae, Metazoa, and Arthropoda, of the query species. The three sets per species excluded either (i) proteins from the query species itself, (ii) all species of the same family or (iii) all species of the same order. The corresponding sets of proteins could provide more or less precise evidence for gene prediction depending on the degree of saturation by closely related species. The level (i) offers the largest number of close relatives while the level (iii) provides the least number of them and the least precise evidence for a query species. We used proteins from the corresponding sets of species selected at the level (iii) for the other eight species.

TSEBRA (with default hyperparameters) had a higher accuracy than either BRAKER1 or BRAKER2 across all 11 species and nearly all test settings, see Table 1. The F1-score of TSEBRA was on average 7.78 percent points higher on gene level, 4.53 percent points higher on transcript level, and 2.06 percent points higher on CDS level than the maximum F1-score of BRAKER1 and BRAKER2. Note that for some species, the BRAKER1 F1–score was higher than the one for BRAKER2 and *vice versa* for other species. The directionality was strongly correlated between the CDS, transcript, and gene levels. For a user, it is difficult to figure out which mode of BRAKER would perform better for a genome of interest. Using TSEBRA is supposed to resolve this uncertainty. TSEBRA generates a higher increase in specificity than in sensitivity: on average Sn increased by 0.52 percent points for all evaluation levels while Sp increased by 8.78 percent points. This was likely caused by the setting of parameters filtering out a majority of transcripts with low support from extrinsic evidence.

**Table 1.** F1-score on CDS, transcript, and gene level for BRAKER1 (RNA-seq hints), BRAKER2 (protein hints of type (iii)), TSEBRA_EVM, EVM using comparable evidence, and TSEBRA (default hyperparameter) with hints generated by the BRAKER runs. For *A. thal*, *C. ele.*, *D. mel.*, a set of genome partitions, each totaling 90% of the genome size, was sampled for the evaluation of all methods. For all other species, the tests were run on the full genomes for BRAKER1, BRAKER2, and TSEBRA. (See Table S1 in Supplementary Information for full species names and Table S2 in Supplementary Information for the results with different protein sets.)

| CDS level F1-score |  |  |  |  |  |
| --- | --- | --- | --- | --- | --- |
|  | BRAKER1 | BRAKER2 | EVM | TSEBRA_EVM | TSEBRA |
| *A. thal.* | 81.87 | 84.01 | 84.41 | 86.21 | **86.90** |
| *B. ter.* | 76.12 | 72.84 |  |  | **77.80** |
| *C. ele.* | 85.87 | 81.13 | **86.14** | 85.13 | 84.48 |
| *D. mel.* | 79.82 | 76.79 | 79.67 | 79.89 | **81.66** |
| *D. rer.* | 74.00 | 72.23 |  |  | **78.40** |
| *M. tru.* | 71.46 | 75.11 |  |  | **80.98** |
| *P. tep.* | **68.61** | 63.90 |  |  | 67.96 |
| *P. tri.* | 78.32 | 83.40 |  |  | **87.60** |
| *R. pro.* | 53.54 | 54.49 |  |  | **56.30** |
| *T. nig.* | 53.95 | 57.97 |  |  | **58.70** |
| *X. tro.* | 74.96 | 75.89 |  |  | **79.44** |
| Transcript level F1-score |  |  |  |  |  |
|  | BRAKER1 | BRAKER2 | EVM | TSEBRA_EVM | TSEBRA |
| *A. thal.* | 53.78 | 56.63 | 57.32 | 61.35 | **62.00** |
| *B. ter.* | 33.15 | 26.49 |  |  | **35.02** |
| *C. ele.* | 53.30 | 42.71 | 52.76 | 54.46 | **55.94** |
| *D. mel.* | 51.33 | 46.94 | 49.90 | 53.76 | **55.18** |
| *D. rer.* | 24.99 | 22.17 |  |  | **33.43** |
| *M. tru.* | 39.04 | 44.09 |  |  | **51.72** |
| *P. tep.* | 26.14 | 18.04 |  |  | **28.89** |
| *P. tri.* | 47.04 | 55.96 |  |  | **62.31** |
| *R. pro.* | 12.84 | 12.65 |  |  | **15.22** |
| *T. nig.* | 5.74 | 7.93 |  |  | **9.78** |
| *X. tro.* | 22.88 | 23.84 |  |  | **31.83** |
| Gene level F1-score |  |  |  |  |  |
|  | BRAKER1 | BRAKER2 | EVM | TSEBRA_EVM | TSEBRA |
| *A. thal.* | 65.51 | 70.58 | 70.88 | 78.35 | **79.69** |
| *B. ter.* | 38.91 | 32.18 |  |  | **44.71** |
| *C. ele.* | 63.13 | 52.29 | 63.98 | 68.90 | **70.78** |
| *D. mel.* | 64.44 | 61.25 | 64.94 | 71.34 | **73.93** |
| *D. rer.* | 31.49 | 27.37 |  |  | **44.13** |
| *M. tru.* | 40.03 | 44.96 |  |  | **54.05** |
| *P. tep.* | 28.59 | 19.99 |  |  | **33.83** |
| *P. tri.* | 53.11 | 63.88 |  |  | **73.45** |
| *R. pro.* | 13.64 | 12.91 |  |  | **16.21** |
| *T. nig.* | 6.59 | 8.87 |  |  | **11.46** |
| *X. tro.* | 26.40 | 30.58 |  |  | **41.26** |

Our tests showed that a single parameter set is sufficient for TSEBRA working with BRAKER1 and BRAKER2 across all the tested genomes, therefore, a change (training) of the set of parameters for each new genome may not be needed. The number of transcripts per gene selected by TSEBRA was on average 1.07 which is at the same level as BRAKER2, and lower than the average of 1.20 observed for BRAKER1.

### Comparison with EVidenceModeler

We also compared the accuracy of TSEBRA with the accuracy of EVidenceModeler (EVM, commit 68e724e from GitHub [35]) working to combine BRAKER1 and BRAKER2 predictions with heterogeneous extrinsic evidence. Comparison of TSEBRA and EVM was performed for the genomes of the model species *A. thaliana*, *C. elegans*, and *D. melanogaster*.

EVM takes extrinsic evidence in form of spliced alignments from assembled transcripts. This type of evidence is not produced by BRAKER1 utilizing mappings of unassembled reads. To make a comparison between EVM and TSEBRA on the same data, we produced new and comparable extrinsic evidence for EVM (i.e. spliced alignments) and TSEBRA (i.e. intron or start/stop codon hints). We used the protein alignments generated by ProtHint during the BRAKER2 run and assembled spliced alignments from the RNA-seq reads sampled by VARUS. For each locus, we selected the protein alignment produced by ProtHint with the highest DIAMOND [36] score creating a genome-wide set of protein alignments. To produce RNA-seq based hints, we reconstructed transcripts from the RNA-seq reads with Trinity (v2.12.0) [37] and applied PASA (v2.4.1) [38] to assemble and align them. The genomes were partitioned into 400, 000 bp long segments with an overlap of 50, 000 bp between neighboring segments employing the tools provided by EVM. For each partition, EVM was run with transcript and protein evidence from PASA and protein alignments, respectively. TSEBRA was run with the introns from both sets of alignments and the start/stop codons from the protein alignments. We refer to this particular TSEBRA run as TSEBRA_EVM.

EVM requires that a weight is assigned to each of the four input sources. We used 10% of the total number of partitions to search for a good set of weights for EVM and a set of hyperparameters for TSEBRA. We used the remaining partitions to evaluate the accuracy of TSEBRA_EVM. The values of the hyperparameters are available in Supplementary Information Table S3 and Table S4.

We compared the accuracy of TSEBRA with EVM, one of the most cited combiner tools to date. EVM was previously used to combine BRAKER with other predictions [39], to combine BRAKER2 predictions with RNA-seq evidence [40, 41] or other RNA-seq based predictions [42], and even for combining multiple BRAKER predictions [43]. Still, it is not the most suitable task for EVM to create a BRAKER-only combination. The authors of EVM recommend the use of a set of gene predictions, usually more than two, along with extrinsic evidence, because the strength of EVM is in finding consensus among diverse sources. This is in conflict with the fact that there is no direct way for EVM to use the hints generated by BRAKER and that we were looking for a way to combine only two gene predictions. In addition, EVM reports only one transcript per gene, which limits the completeness of its annotation output in a setting with much evidence for alternative splicing.

We compared TSEBRA and EVM to address a question: which is the better method for combining BRAKER1 and BRAKER2 predictions? In a test setting with comparable extrinsic evidence, we evaluated them across three species with three different protein sets each. Both methods have successfully combined the BRAKER1 and BRAKER2 predictions into one set with increased F1-score, see Figure 3. Still, TSEBRA_EVM had, compared to EVM, a higher accuracy on average with an average increase of the F1-score at the gene, transcript, and CDS levels of 6.12, 3.38, and 0.79 percent points, respectively. These improvements came with an overall increased Sn and Sp for TSEBRA_EVM on the transcript and gene levels. Only at the CDS level, both methods had a similar accuracy; EVM had a slightly higher Sn and TSEBRA_EVM had a higher Sp.

**Figure 3.**
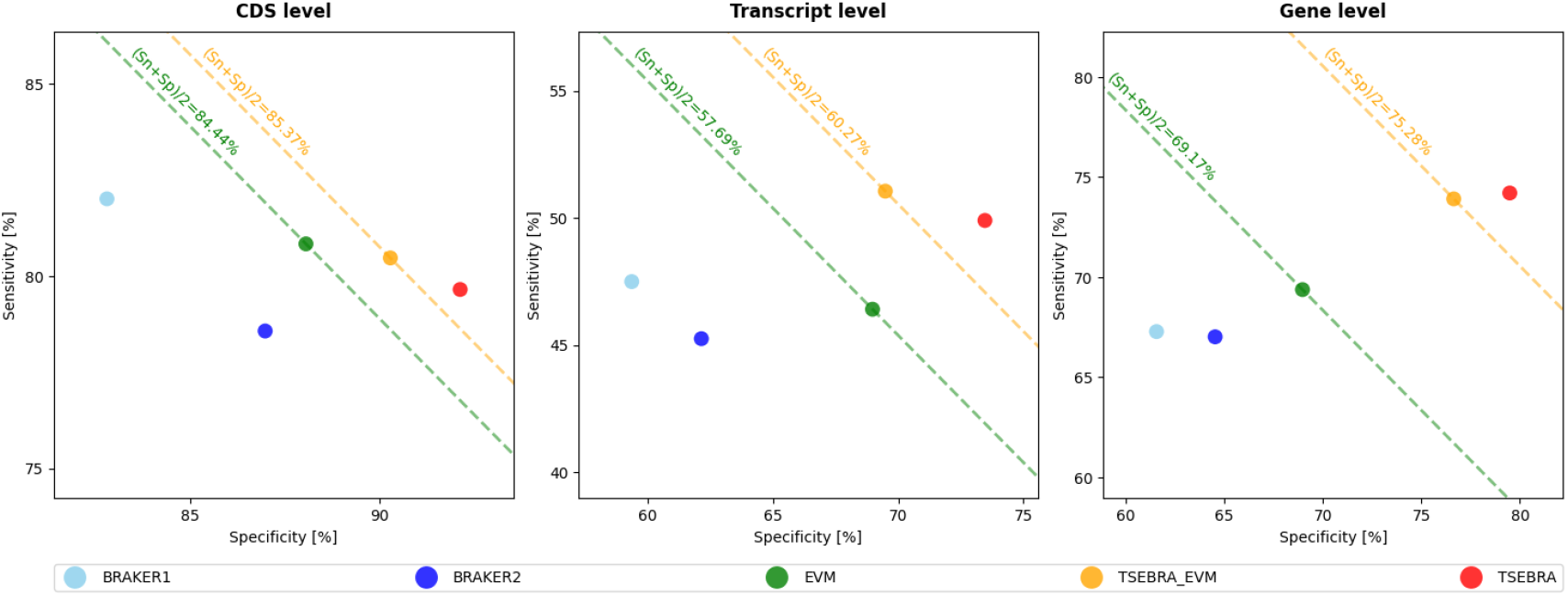
Average gene, transcript, and CDS levels Sn and Sp for all nine combinations of three model species (*A. thal*, *C. ele.*, *D. mel.*) and all test settings ((i), (ii), (iii)) as described for these model species in Table 1.

We had to carefully select the test setting since the choice of partition size made a difference to EVM when it makes models of intergenic regions. An additional difficulty was that the tools used for creating the partitions enforced hard borders, even if they split a transcript. To neutralize this issue, an overlap between partitions was used.

TSEBRA executes much faster than EVM, which constructs a complete gene model and evaluates all possible gene structures. TSEBRA, on the other hand, only evaluates the set of explicitly given transcripts as input. In our tests, we ran both methods separately in parallel on a 28 multi-core processor, the average runtime of EVM was 35.28 min and of TSEBRA_EVM 0.37 min.

## Conclusions

We presented TSEBRA, a tool that selects more reliable gene predictions (transcripts) from the sets of transcripts generated independently by BRAKER1 and BRAKER2. A novel approach to transcript selection was successfully implemented. In computational experiments made with genomes of 11 diverse eukaryotic species we have shown that the set of transcripts selected by TSEBRA matched annotated genes (believed to be the true ones) with higher accuracy than both BRAKER1 and BRAKER2. Note that the combined extrinsic evidence is not used at the step of generation of gene predictions. BRAKER1 and BRAKER2 use disjoint evidence sources also for *training* statistical gene-finding models. A relative complementarity of the gene sets can be an advantage when they are combined subsequently. The ranking and selection of the final set of transcripts, however, does use both protein and RNA-seq evidence. This approach makes an effective use of both sources of extrinsic evidence for selection of most likely true positive transcripts from the set of candidates, the transcripts generated by BRAKER1 and BRAKER2 running in parallel.

Thus, TSEBRA makes a useful tool that with help of heterogeneous extrinsic evidence transforms the union of predictions of BRAKER1 and BRAKER2 into a set of gene predictions whose accuracy exceeds the accuracy of both BRAKER1 and BRAKER2 running separately.

## Availability and requirements

Project name: TSEBRA.

Project home page: https://github.com/Gaius-Augustus/TSEBRA.

Operating system(s): Linux, MacOS.

Programming language: Python.

Other requirements: Python 3.0 or higher.

License: Artistic License 2.0 (see https://opensource.org/licenses/Artistic-2.0).

Any restrictions to use by non-academics: Artistic License 2.0 restrictions apply.

## Abbreviations

NCBI: National Center for Biotechnology Information
RNA-seq: RNA-sequencing
SRA: Sequence Read Archive
EVM: EVidenceModeler
TSEBRA: Transcript Selector for BRAKER
Sp: specificity
Sn: sensitivity

## Supplementary Information

**Additional file 1.pdf**. Supplementary tables (S1-S4).

## Availability of data and materials

Archived source code of TSEBRA as at time of publication: https://github.com/Gaius-Augustus/TSEBRA/releases/tag/v1.0.1. Description of how to generate the results of the experiments: https://github.com/Gaius-Augustus/TSEBRA-experiments.

## Competing interests

The authors declare that they have no competing interests.

## Authors’ contributions

MS and LG designed TSEBRA. LG implemented TSEBRA, ran BRAKER1, EVM and performed all accuracy comparisons. TB ran BRAKER2. MB supervised BRAKER2 experiments. TB and KJH sampled reads with VARUS. LG, KJH, MB, and MS participated in writing the manuscript. All authors read and approved the manuscript.

## Funding

The research was supported in part by the US National Institutes of Health grant GM128145.

## Supporting Information

If you intend to keep supporting files separately you can do so and just provide figure captions here. Optionally make clicky links to the online file using `\href{url}{description}`.

## Notes

https://github.com/Gaius-Augustus/TSEBRA

### Competing Interest Statement

The authors have declared no competing interest.

