## Supplementary tables (S1-S4) for "TSEBRA: Transcript Selector for BRAKER"

**Table S1** List of all genomes and annotation versions used for the experiments. The species with * are the model species used for the comparison of TSEBRA and EVM.

| Species | Annotation version | Genome size (Mb) | Number of genes |
|---|---|---|---|
| *Arabidopsis thaliana** | Tair Araport 11 (Jun 2016) | 119 | 27,445 |
| *Bombus terrestris* | NCBI Annotation Release 102(Apr 2017) | 249 | 10,581 |
| *Caenorhabditis elegans** | WormBase WS271 (May 2019) | 100 | 20,172 |
| *Danio rerio* | Ensembl GRCz11.96 (May 2019) | 1345 | 25,254 |
| *Drosophila melanogaster** | FlyBase R6.28 (Jun 2019) | 138 | 13,929 |
| *Medicago truncatula* | MtrunA17r5.0-ANR-EGN-r1.6 (Feb 2019) | 430 | 44,464 |
| *Parasteatoda tepidariorum* | NCBI Annotation Release 101 (May 2017) | 1445 | 18,602 |
| *Populus trichocarpa* | JGI Ptrichocarpa 533 v4.1 (Nov2019) | 389 | 34,488 |
| *Rhodnius prolixus* | VectorBase RproC3.3 (Oct 2017) | 707 | 15,061 |
| *Tetraodon nigroviridis* | TETRAODON8.99 (Nov 2019) | 359 | 19,589 |
| *Xenopus tropicalis* | NCBI Annotation Release 104 (Apr 2019) | 1449 | 21,821 |

**Table S2** F1-score on CDS, transcript, and gene level for BRAKER1 (RNA-seq hints), BRAKER2 (protein hints), TSEBRA_EVM, EVM using comparable evidence, and TSEBRA with hints generated by the BRAKER runs. The methods were run three times for each species, allowing different minimum evolutionary distances ((i) species excluded, (ii) family excluded or (iii) order excluded) for the proteins used as support. A different set of genome partitions, each totaling 90% of the genome size from a species, was sampled for the evaluation of all methods for each model species and test setting.

| **CDS level F1-score** | | | | | |
|---|---|---|---|---|---|
| | BRAKER1 | BRAKER2 | EVM | TSEBRA_EVM | TSEBRA |
| *A. tha.*(i) | 81.67 | 84.62 | 85.10 | **87.70** | 87.34 |
| *A. tha.*(ii) | 81.67 | 84.00 | 84.07 | 86.46 | **86.93** |
| *A. tha.*(iii) | 81.87 | 84.01 | 84.41 | 86.21 | **86.90** |
| *C. ele.*(i) | 85.76 | 87.26 | 88.15 | 88.03 | **88.27** |
| *C. ele.*(ii) | 85.83 | 80.91 | **86.46** | 85.05 | 84.46 |
| *C. ele.*(iii) | 85.87 | 81.13 | **86.14** | 85.13 | 84.48 |
| *D. mel.*(i) | 79.88 | 84.03 | 83.74 | 85.37 | **85.71** |
| *D. mel.*(ii) | 79.20 | 79.86 | 80.78 | 81.81 | **82.95** |
| *D. mel.*(iii) | 79.82 | 76.79 | 79.67 | 79.89 | **81.66** |
| **Transcript level F1-score** | | | | | |
| | BRAKER1 | BRAKER2 | EVM | TSEBRA_EVM | TSEBRA |
| *A. tha.*(i) | 53.52 | 60.25 | 61.01 | **64.92** | 64.05 |
| *A. tha.*(ii) | 53.48 | 57.02 | 56.63 | 61.97 | **62.14** |
| *A. tha.*(iii) | 53.78 | 56.63 | 57.32 | 61.35 | **62.00** |
| *C. ele.*(i) | 53.44 | 55.13 | 57.41 | 59.66 | **60.54** |
| *C. ele.*(ii) | 53.36 | 42.52 | 53.79 | 54.58 | **56.09** |
| *C. ele.*(iii) | 53.30 | 42.71 | 52.76 | 54.46 | **55.94** |
| *D. mel.*(i) | 51.44 | 58.50 | 57.86 | **62.26** | 61.61 |
| *D. mel.*(ii) | 51.22 | 50.98 | 52.23 | 56.37 | **57.01** |
| *D. mel.*(iii) | 51.33 | 46.94 | 49.90 | 53.76 | **55.18** |
| **Gene level F1-score** | | | | | |
| | BRAKER1 | BRAKER2 | EVM | TSEBRA_EVM | TSEBRA |
| *A. tha.*(i) | 65.20 | 75.03 | 75.09 | **81.45** | 81.15 |
| *A. tha.*(ii) | 65.22 | 71.08 | 70.07 | 78.92 | **79.77** |
| *A. tha.*(iii) | 65.51 | 70.58 | 70.88 | 78.35 | **79.69** |
| *C. ele.*(i) | 63.25 | 67.18 | 69.37 | 73.80 | **76.31** |
| *C. ele.*(ii) | 63.12 | 52.15 | 65.18 | 69.30 | **70.92** |
| *C. ele.*(iii) | 63.13 | 52.29 | 63.98 | 68.90 | **70.78** |
| *D. mel.*(i) | 64.56 | 75.31 | 74.75 | 80.66 | **81.46** |
| *D. mel.*(ii) | 64.12 | 66.08 | 67.81 | 74.46 | **76.07** |
| *D. mel.*(iii) | 64.44 | 61.25 | 64.94 | 71.34 | **73.93** |

**Table S3** Weights for each input source used by the EvidenceModeler for the comparison with TSEBRA_EVM for each model species and all test settings.

| | BRAKER | BRAKER2 | RNA-seq | Protein |
|---|---|---|---|---|
| *A. tha.*(i) | 30 | 6750 | 22 | 22 |
| *A. tha.*(ii) | 30 | 6750 | 7 | 6750 |
| *A. tha.*(iii) | 30 | 6750 | 22 | 22 |
| *C. ele.*(i) | 30 | 480 | 900 | 480 |
| *C. ele.*(ii) | 30 | 22 | 60 | 7 |
| *C. ele.*(iii) | 30 | 22 | 60 | 22 |
| *D. mel.*(i) | 30 | 6750 | 45 | 6750 |
| *D. mel.*(ii) | 30 | 480 | 6750 | 0 |
| *D. mel.*(iii) | 30 | 22 | 60 | 30 |

**Table S4** Hyperparameters used by TSEBRA_EVM for the comparison with TSEBRA_EVM for each model species and all test settings. The hyperparameter sets are described in Implementation.

| | **Evidence source weight** | | **Low Evidence support threshold** | | **Transcript score threshold** | | | |
|---|---|---|---|---|---|---|---|---|
| | Protein ($w_P$) | RNA-seq ($w_R$) | Intron | Start/stop-codon | $\epsilon_1$ | $\epsilon_2$ | $\epsilon_3$ | $\epsilon_4$ |
| *A. tha.*(i) | 15.0 | 10.0 | 0.75 | 1.0 | 0.0 | 0.0 | 20.0 | 0.0 |
| *A. tha.*(ii) | 10.0 | 0.5 | 0.75 | 1.0 | 0.0 | 0.0 | 10.0 | 0.0 |
| *A. tha.*(iii) | 5.5 | 10.0 | 0.5 | 1.0 | 0.0 | 0.5 | 30.0 | 2.5 |
| *C. ele.*(i) | 5.5 | 15.0 | 0.0 | 0.0 | 0.0 | 1.0 | 10.0 | 2.5 |
| *C. ele.*(ii) | 0.5 | 10.0 | 0.0 | 0.0 | 0.0 | 0.5 | 20.0 | 10.0 |
| *C. ele.*(iii) | 0.5 | 10.0 | 0.0 | 0.0 | 0.0 | 0.5 | 10.0 | 15.0 |
| *D. mel.*(i) | 1.0 | 0.5 | 0.875 | 0.5 | 0.0 | 0.0 | 5.0 | 0.0 |
| *D. mel.*(ii) | 5.5 | 0.75 | 0.25 | 1.0 | 0.0 | 0.5 | 20.0 | 10.0 |
| *D. mel.*(iii) | 5.5 | 1.0 | 0.25 | 0.5 | 0.0 | 0.5 | 15.0 | 15.0 |
